# Backbone Rigidity Encodes Universal Viscoelastic Signatures in Biomolecular Condensates

**DOI:** 10.1101/2025.06.10.658939

**Authors:** Sean Yang, Subhadip Biswas, Davit A Potoyan

**Affiliations:** Department of Chemistry, Iowa State University, Ames IA 50011, USA; Roy J. Carver Department of Biochemistry, Biophysics, and Molecular Biology, Ames IA 50011, USA

## Abstract

Biomolecular condensates exhibit a wide range of viscoelastic properties shaped by their molecular sequence and composition. Coarse-grained molecular models of intrinsically disordered proteins are widely used to complement experiments by revealing the structure and thermodynamics of condensates. However, fully flexible chain representations of inherently disordered proteins often fail to capture their complex viscoelastic behavior, instead predicting purely viscous responses. In this work, we demonstrate that introducing sequence-dependent chain rigidity enables the accurate reproduction of the elastic and viscous moduli for experimentally characterized condensates of A1-LCD and its numerous mutants. Furthermore, we show that the frequency-dependent loss factor can be described by a single parameter that universally correlates with viscosity across different sequences and variations of the coarse-grained molecular energy function. Our results also reveal that increased chain rigidity, indicated by a larger gyration radius, expands the condensates’ elastic regime. Finally, we elucidate the microscopic origins of sequence-encoded viscoelasticity by showing how it can be tuned through sequence rearrangements that promote sticker cluster formation.

## I. INTRODUCTION

The phase separation of biomolecules into membraneless condensates is a ubiquitous organizing mechanism in cells for exerting spatial and temporal control over biochemical processes [1–4]. The functional attributes of condensates stem from their liquid-like properties, which manifest in their ability to swiftly assemble, disassemble, fuse, and divide, allowing cells to rapidly adapt to changing environmental cues [5– 8]. Over time, however, condensates may lose their liquid-like material properties and age into more solid-like assemblies associated with pathological conditions [9–12]. Depending on their molecular composition, condensates span a wide range of material properties: from Newtonian fluids to viscoelastic materials and gel-like states, reflecting their capacity to integrate, process, and store regulatory information at the mesoscale [13–18].

Capturing this rich spectrum of material behaviors in silico remains a significant challenge, particularly for computational models aiming to bridge molecular sequence features with mesoscopic rheology. Microrheology experiments have been instrumental in uncovering the sequence- and composition-dependent viscoelastic properties of biomolecular condensates [19–21]. In addition to measuring material responses, microrheology provides access to internal dynamics and characteristic timescales associated with aging, relaxation, and flow [10, 22, 23]. Complementing these experimental advances, sequence-resolved coarse-grained models have played a central role in elucidating the molecular origins of condensate behavior. Notably, such models have successfully reproduced key thermodynamic trends of intrinsically disordered proteins, including saturation concentrations and critical temperatures [18, 24–28]. A key assumption in many single-residue, sequence-resolved models is that chains are fully flexible, an approach that significantly accelerates simulations, prevents entanglement, and enables robust predictions of structural and thermodynamic properties [29]. Driven by available experimental data from SAXS and turbidity measurements, most molecular models prioritize thermodynamic accuracy and structural realism over dynamic material properties, thereby overlooking the subtle role of local chain rigidity in shaping stress relaxation and energy dissipation. Recent reports [30, 31] have shown that fully flexible chains fail to capture viscoelastic crossover behavior when applied to material properties. It has been well characterized that chains of intrinsically disordered proteins will have sequence-dependent rigidity, whereby flexible glycine repeats make parts of the chain more flexible while more bulky hydrophobic and aromatic residues often lead to a non-linear effect on rigidity, leading to deviations from polymer scaling behaviour [32– 34]. Single-molecule and simulation studies have also revealed that intrinsically disordered proteins exhibit sequence-dependent internal friction, reflecting barriers to conformational rearrangement even in the absence of crowding [35– 37]. While such internal friction contributes to the intrinsic timescales of chain reconfiguration, condensates also exhibit additional collective viscoelastic effects arising from intermolecular interactions, transient crosslinking, and network formation [19, 31, 38]. In this work, we demonstrate that introducing semiflexibility into the C_α_ coarse-grained molecular models of A1-LCD enables the reproduction of experimental viscoelastic trends [39]. Furthermore, we systematically investigate how sequence-dependent variations in local chain rigidity affect the behavior of condensates. Our findings demonstrate that incorporating sequence-dependent rigidity enables accurate prediction of experimentally observed rheological trends in hnRNP A1, for which detailed microrheology data are available [39]. By designing new sequences via reshuffling of A1 LCD residues, we also provide experimentally testable predictions for tuning the viscoelasticity of condensates by systematic variation of hydrophobic patterns. We elucidate the microscopic origins of sequence-encoded viscoelasticity by showing how it can be tuned through sequence rearrangements that promote sticker cluster formation. Finally, we show that a single master curve can capture viscoelastic trends by appropriately rescaling viscous and elastic moduli.

This reflects that time scales dictating material properties are captured by sequence-dependent chain rigidity and the onset of the collapse transition. Our results establish a predictive framework for engineering the material properties of condensates via sequence design, preparing the ground for rational modulation of condensate functional dynamics in cells.

## II. METHODS

### Description of models

We have employed a one-bead-per-residue hydrophobicity scale (HPS) [29] models with parameter CALVADOS 2 [40] to simulate the low-complexity domain of protein hn-RNP A1 (referred to as A1-LCD in the following text) [39]. The coarse-grained (CG) beads in the chain represent the amino acids of peptides. The non-bonded interactions between residues *i, j* are modeled via the Ashbaugh-Hatch potential, and salt-screened electrostatic interactions are modeled via the Debye-Huckel potential:

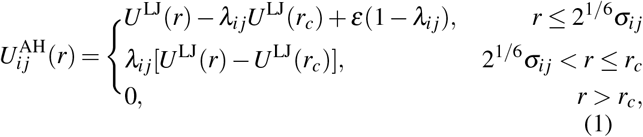

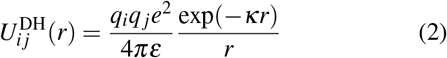

where *ε* = 0.8368 kJ/mol, *r*_*c*_ = 4 nm, and *U* ^LJ^ is the Lennard-Jones potential *U* ^LJ^(*r*) = 4*ε* (*σ*_*i j*_*/r*)^12^ − (*σ*_*i j*_*/r*)^6^. *σ*_*i j*_ = (*σ*_*i*_ + *σ*_*j*_)*/*2 and *λ*_*i j*_ = (*λ*_*i*_ + *λ*_*j*_)*/*2 are arithmetic averages of monomer size and hydrophobicity value, respectively. The pair potential with *λ* = 0 consists of only the repulsive term equivalent to the Weeks-Chandler-Andersen functional form [41]. We use *σ* values of amino acids from van der Waals volumes [42]. We use *λ* values of amino acids from the recently proposed CALVADOS 2 parameters [40]. In the Debye-Huckel potential, *q* is the charge number of particles, and 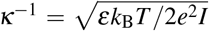 is the Debye length, where *ε* is the dielectric constant and *I* is the ionic strength. We take *I* = 0.17M, *T* = 300 K and *κ*^−1^ = 0.9nm in our simulations. The temperature-dependent dielectric constant used in our simulations has the following empirical relation: *ε*(*T*)*/ε*_0_ = 5321*/T* 233.76 − 0.9297*T* 1.41710 *×* 10^−3^*T* ^2^ − 8.29210 *×* 10^−7^*T* ^3^. All coarse-grained beads are connected via bonded interactions modeled by harmonic potentials.

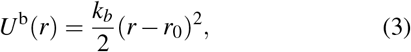

where *k*_b_ = 8033 kJ/mol/nm^2^ and *r*_0_ = 0.38 nm. We employed a harmonic angular potential between two adjacent bonds in the simulations. The angular potential of residue *i* (Fig. 1A) is expressed as

**FIG. 1.**
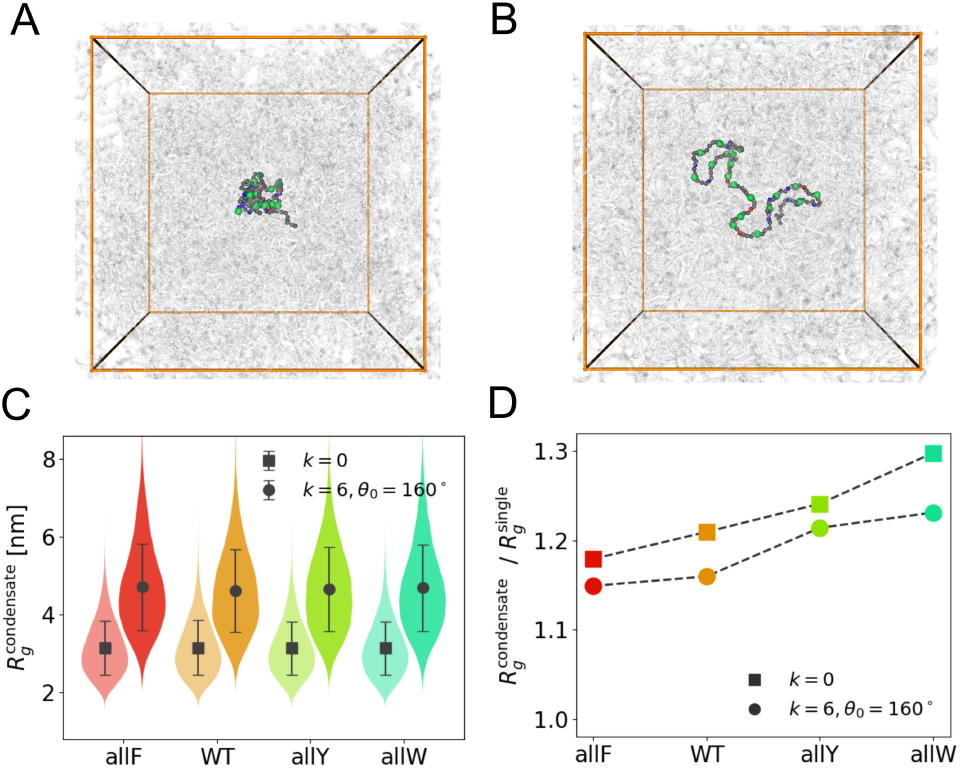
Chain rigidity and conformation. **(A)** Snapshot of a WT A1-LCD in a condensate environment with *k* = 0. **(B)** Snapshot of a WT A1-LCD peptide in a condensate environment with *k* = 6 *k*_*B*_*T* /rad^2^ and *θ*_0_ = 160^°^. **(C)** Radius of gyration *R*_*g*_ for four mutants of A1-LCD (allF, WT, allY and allW) under two different *U*^*a*^ parameter sets represented by different symbols. Squares and circles indicate *k* = 0 and *k* = 6 *k*_*B*_*T* /rad^2^, *θ*_0_ = 160^°^ in condensate simulations. **(D)** Ratios of *R*_*g*_ between condensates and single-chain simulations for mutants under the same *U*^*a*^ parameter sets as in C.

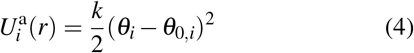

where *k* the torsional rigidity is in units *k*_*B*_*T* /rad^2^ and *θ*_0,*i*_ is the natural angle between two bonds of the residue *i*. These are adjustable parameters, and their variation is systematically explored in the Results section.

### Molecular simulations and viscoelasticity measurements

All simulations were performed using the OpenMM (v7.7) molecular dynamics library on GPU nodes. The number of chains in the simulation box was *N*_pep_ = 250. The initial length of the cubic box was set at 50 nm, much larger than the equilibrium size, to make a relatively low density of the chain for initialization. After energy minimization, we used the *NVT* ensemble with a timestep of 0.01 ps. Next, by gradually shrinking the box dimensions, we slowly increase the density of our system until it reaches the condensate density in previous phase separation simulations. The parameters of this process are shown in Supplementary Information Section A (SI-A). Finally, we fix the box size and carry out 1 *×* 10^8^step *NVT* simulations for all the following measurements.

By the Green-Kubo (GK) relation, the shear stress relaxation modulus *G*(*t*) can be determined by computing the autocorrelation of the off-diagonal components of the stress tensor. A more accurate expression can be obtained by taking into account the isotropic nature of the system [43, 44]

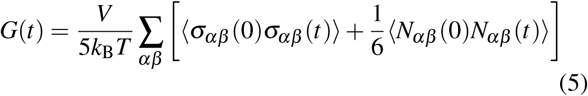

where *N*_*αβ*_ = *σ*_*αα*_ − *σ*_*ββ*_ is the normal stress difference and the summation contains (*αβ* = *xy, yz, xz*). To avoid the typical noisy nature of *G*(*t*) at a long time scales, we fitted *G*(*t*) to a series of Maxwell modes *G*_M_(*t*) = S_*m*_*G*_*m*_exp(− *t/τ*_*m*_) which are equidistant in logarithmic time [44, 45]. Then, elastic and viscous moduli *G*^*′*^(*ω*) and *G*^*′′*^(*ω*) can be straightforwardly calculated by the Fourier transform of *G*(*t*). The shear viscosity can be calculated by integrating the shear stress relaxation modulus over the relaxation time.

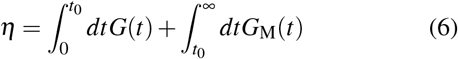

The division time *t*_0_ is chosen so all the fast intra-molecular oscillations of *G*(*t*) have decayed, and the function becomes positive after *t*_0_. We recorded the stress tensor matrix every 20 steps for evaluating the short-time integral and 200 steps for assessing the long-time integral of Eq. (6) in the *NVT* simulations.

## III. RESULTS

We systematically modulate local chain rigidity in the HPS model by adjusting the angular potential *U*^*a*^.

### Chain conformation in dilute and dense phases

Before analyzing collective viscoelasticity and the role of chain rigidity in network-level relaxation, it is instructive to examine how it affects the conformational properties of individual chains, both inside and outside the condensate environment. The wild-type (WT) A1-LCD undergoes significant extension upon the application of a harmonic angular potential (Fig. 1A-B). This extension is quantified by the radius of gyration *R*_*g*_ (Fig. 1C), which increases with local backbone rigidity, as expected from the imposed angular constraints. In the condensed phase, chain extension is further enhanced relative to the dilute phase, as indicated by a ratio 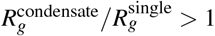 (Fig. 1D). This effect is consistent with prior observations that the sticky environment of condensates, both at their interfaces and interiors, can promote chain expansion [25, 46].

We examined A1-LCD variants with systematic substitutions of aromatic residues to assess how hydrophobicity influences these conformational trends. Based on hydrophobic interaction strength (F *<* Y *<* W), the net inter-residue attraction follows the order allF *<* WT *<* allY *<* allW (see SI-A for sequences). In single-chain simulations, increasing hydrophobicity leads to a modest reduction in *R*_*g*_, reflecting a higher likelihood of intramolecular contacts among distant residues. This effect is captured by the increasing ratio 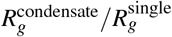 across the series (Fig. 1D).

Interestingly, in condensate simulations, *R*_*g*_ remains largely unchanged across all mutants (Fig. 1C), indicating that enhanced inter-chain attractions do not result in further chain compaction. This reinforces the view that intra-chain attractions primarily govern compaction in dilute conditions, while condensate environments allow for more extended conformations.

### Modulation of viscoelastic moduli

We first examine how the viscoelastic moduli respond to variations in torsional rigidity *k* and the angular potential’s preferred angle *θ*_0_. For weak angular potentials, peptide condensates exhibit dominantly viscous behavior, where the loss modulus *G*^*′′*^(*ω*) exceeds the storage modulus *G*^*′*^(*ω*) across all frequencies. As shown in Fig. 2A-B, this viscous-dominated regime persists for *k* = 0 across all aromatic variants. As *k* increases, local chain rigidity strengthens and the chains straighten. This promotes elastic response: *G*^*′*^(*ω*) grows more rapidly than *G*^*′′*^(*ω*), leading to a crossover into a dominantly elastic regime at intermediate frequencies. For example, at *k* = 6 in the allW sequence, *G*^*′*^ exceeds *G*^*′′*^ near 10^−4^ ps^−1^ (Fig. 2C). The first crossover frequency, *ω*_*R*_ = 2.2 *×* 10^−4^ ps^−1^, corresponds to a characteristic relaxation time *τ*_*R*_ *≡* 1*/ω*_*R*_ = 4.5 ns (Fig. 2D). When rigidity is further increased to *k* = 20, *τ*_*R*_ for the allW sequence rises to 19 ns (Fig. 2E-F), indicating slower relaxation due to enhanced stiffness. A similar transition from viscous to elastic behavior is observed by increasing *θ*_0_ from 140^°^ to 160^°^ at fixed *k* = 10 (Fig. S2C of SI-C).

**FIG. 2.**
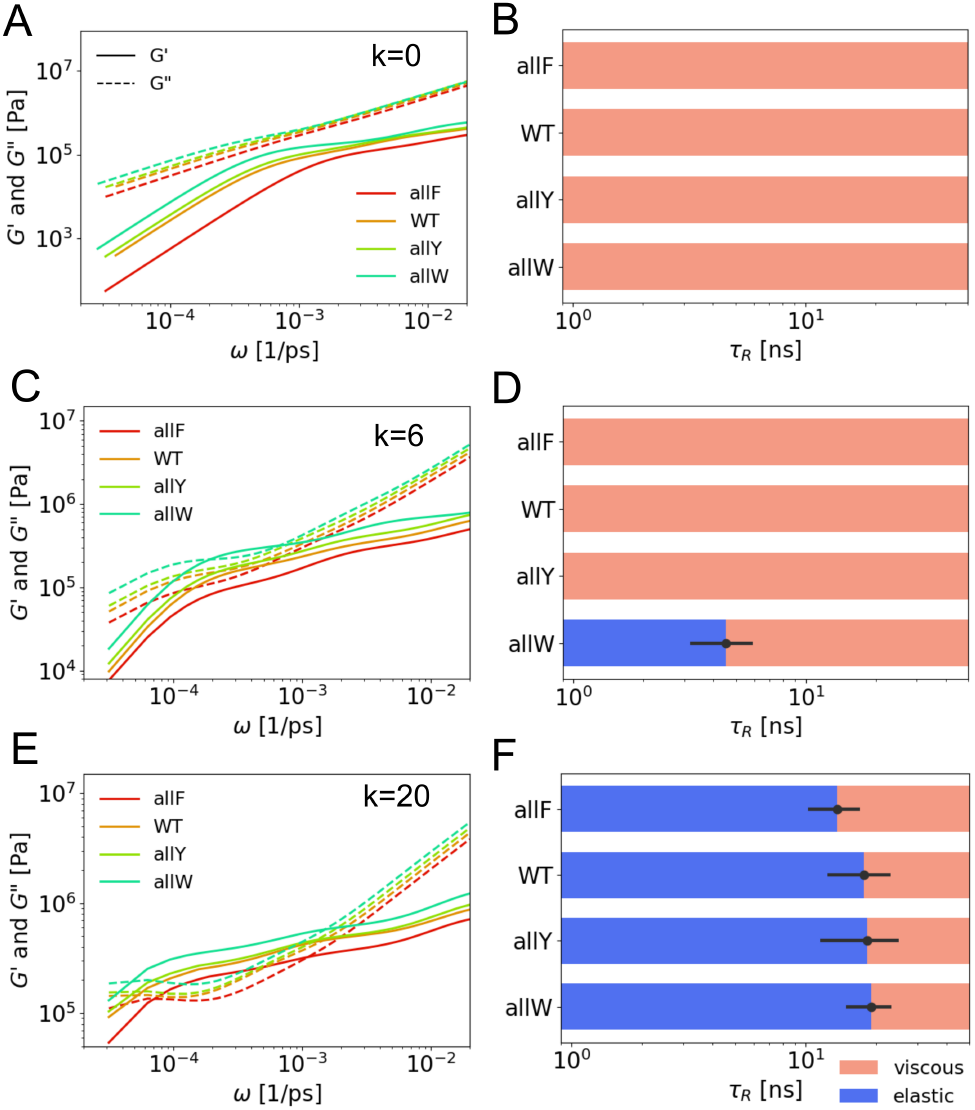
Angular potential modulates viscoelastic moduli. **(A)** Elastic *G*^*′*^(*ω*) and viscous *G*^*′′*^(*ω*) moduli measured from four A1-LCD mutants simulations at *k* = 0. **(B)** Relaxation times *τ*_*R*_ of corresponding crossover points in A. **(C)** *G*^*′*^(*ω*), *G*^*′′*^(*ω*) for four aromatic mutants at *k* = 6 *k*_*B*_*T* /rad^2^ and *θ*_0_ = 160^°^. **(D)** Relaxation times *τ*_*R*_ of corresponding crossover points in C. **(E)** *G*^*′*^(*ω*), *G*^*′′*^(*ω*) for four aromatic mutants at *k* = 20 *k*_*B*_*T* /rad^2^ and *θ*_0_ = 160^°^. **(F)** Relaxation times *τ*_*R*_ of corresponding crossover points in E.

We next investigate how the viscoelastic response depends on aromatic mutations of A1-LCD. In the absence of angular rigidity, increasing hydrophobic attraction among aromatic residues (from allF to allW) raises both *G*^*′*^ and *G*^*′′*^, with *G*^*′*^ rising at a faster rate. As a result, the loss factor *G*^*′′*^*/G*^*′*^ decreases but remains above 1 (Fig. 2A), indicating a viscous-dominated regime persists despite increased interaction strength. Upon introducing moderate angular rigidity (*k* = 6), a transition to a dominantly elastic regime (*G*^*′′*^*/G*^*′*^ *<* 1) is observed across the mutant series, particularly for sequences with stronger hydrophobic interactions. However, the resulting relaxation times *τ*_*R*_ (Fig. 2C-D) remain significantly smaller than experimental values, indicating that this level of rigidity is insufficient to capture the observed viscoelastic timescales fully. At high rigidity (*k* = 20), an elastic-dominated regime is present across all sequences, and *τ*_*R*_ increases systematically from allF to allW (Fig. 2E-F).

As local rigidity and inter-chain hydrophobic interactions increase, the effective viscosity *η* also rises, reflecting stronger network connectivity and potential entanglement. This trend is consistent with the microscopic mechanisms of viscosity [38, 47, 48], as previously analyzed in our recent work. The corresponding viscosity values for each simulation condition are provided in Fig. S2A of SI-C.

### Residue-dependent angular potential

To capture experimentally observed trends in A1-LCD sequence variants [39], we introduce a residue-dependent angular potential 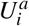 to modulate local chain rigidity based on amino acid identity. Due to their bulky side chains, aromatic residues naturally increase the bond angles of adjacent peptide bonds. To model this effect, we assign a preferred bond angle *θ*_Ar_ for aromatic residues (F, Y, W), with increasing stiffness: *θ*_F_ = 145^°^, *θ*_Y_ = 160^°^, and *θ*_W_ = 175^°^.

The preferred bond angle at position *i* is also modulated by aromatic residues at adjacent positions *i ±* 1. We implement three angular schemes (I, II, III) to test different coupling rules between residue identity and local rigidity (Fig. 3A):

**FIG. 3.**
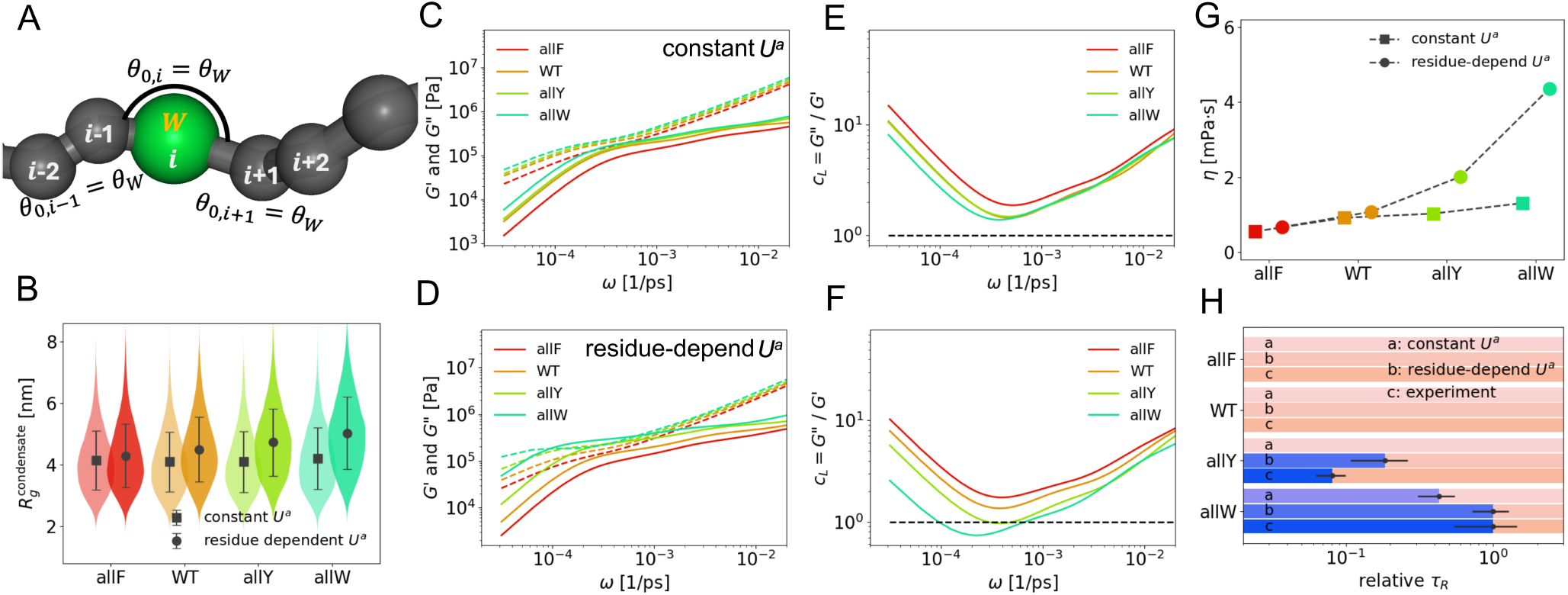
Effect of residue-dependent angular potential. **(A)** Illustration of the residue dependent natural angle *θ*_0,*i*_ in angular potential. As an example of setting III, when the (*i*)-th residue is aromatic (W), the corresponding angles are *θ*_0,*i* 1_ = *θ*_*W*_, *θ*_0,*i*_ = *θ*_*W*_ and *θ*_0,*i*+01_ = *θ*_*W*_. **(B)** Radius of gyration *R*_*g*_ measured from condensate simulations of four mutants under two *U*^*a*^ parameter sets, which is represented by squares (*k* = 16, *θ*_0_ = 140^°^) and circles (residue-dependent setting III). The details of parameter sets are provided in Eq.(7). **(C-F)** *G*^*′*^(*ω*), *G*^*′′*^(*ω*) and loss factor *c*_*L*_ = *G*^*′′*^*/G*^*′*^ measured from simulations of four aromatic mutants using the constant *U*^*a*^ (*k* = 16, *θ*_0_ = 140^°^) and residue-dependent parameter setting III. **(G)** Viscosity *η* measured from condensate simulations of four mutants under two *U*^*a*^ parameter sets, the same as B. **(H)** Delineation of the dominantly viscous (pink) versus elastic (purple) regimes in terms of relaxation times *τ*_*R*_. For each sequence (allF, WT, allY and allW), *τ*_*R*_ is compared among simulations with constant *U*^*a*^ (*k* = 6, *θ*_0_ = 160^°^), residue-dependent *U*^*a*^ (*k* = 16, *θ*_0_ = 140^°^ with setting III), and the experiment [39]. Both simulation and experimental results are normalized by their maximum *τ*_*R*_ of all mutants.

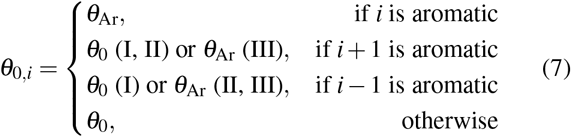

Simulations were conducted using four parameter settings, all with *k* = 16 and baseline *θ*_0_ = 140^°^, to compare uniform and residue-dependent angular potentials. The constant *U*^*a*^ case serves as a reference for isolating the effect of sequence-dependent rigidity (see Table 1 and Eq. 7). These settings progressively increase local rigidity in regions enriched with aromatic residues.

We first analyze changes in chain conformation. For a constant angular potential, the radius of gyration (*R*_*g*_) remains similar across all mutants. However, with residue-dependent angular potentials, *R*_*g*_ increases as the side chain bulk increases from allF to allW (Fig. 3B), indicating enhanced chain extension due to local rigidity amplification. Next, we examine how viscoelastic moduli respond. Both *G*^*′*^ and *G*^*′′*^ increase when residue-specific rigidity is introduced, compared to the constant angular potential case (Fig. 3C-D). This trend mirrors the effect of increasing *k* uniformly (Fig. 2).

The residue-dependent angular potential also amplifies differences in viscoelasticity across mutants. As shown in Fig. 3E-F, the loss factor *c*_*L*_(*ω*) *≡ G*^*′′*^*/G*^*′*^ exhibits greater spread among mutants than under a constant potential. This enhanced contrast is also reflected in the viscosity *η* values across mutants (Fig. 3G). At low frequencies, we observe the characteristic terminal relaxation scaling: *G*^*′*^(*ω*) ∝ *ω*^2^ and *G*^*′′*^(*ω*) ∝ *ω*, implying *c*_*L*_(*ω*) ∝ *ω*^−1^ [17]. In the intermediate-frequency regime, the slower increase of *G*^*′*^(*ω*) produces a turning point in each *c*_*L*_(*ω*) curve (Fig. 3E).

To quantify the transition between viscous and elastic behavior, we extract the relaxation time *τ*_*R*_ from the crossover frequency (*G*^*′*^ = *G*^*′′*^), where present (Fig. 3H). In cases with no crossover, no *τ*_*R*_ is reported. We compare three conditions: (1) simulations with a constant angular potential (*k* = 6, *θ*_0_ = 160^°^), (2) simulations with residue-dependent rigidity (*k* = 16, *θ*_0_ = 140^°^, scheme II), and (3) experimental data from [39]. For comparison, *τ*_*R*_ is normalized by the maximum value observed for the allW mutant in each dataset. The results demonstrate that simulations incorporating residue-dependent angular potentials yield relaxation times that more closely match experimental trends than models with constant rigidity, highlighting the importance of sequence-encoded stiffness in capturing viscoelastic behavior.

### Tuning viscoelasticity of condensates via rearranging sequences

To provide experimentally testable predictions regarding the effects of chain rigidity modulation, we analyzed the rearrangement of representative sequences (allF, WT, allY, and allW). In rearrangement type I, aromatic residues are clustered in the center of the chain; in rearrangement type II, they are also centered but spaced with one-residue gaps. The exact rearranged sequences are provided in SI-A.

Upon rearrangement type II, the *R*_*g*_ of the allW sequence increases by approximately 20%, whereas the allF sequence exhibits only a modest change (5%) (Fig. 4A-B). This trend is consistently observed in both single-chain and condensate simulations, indicating that the larger side chain in tryptophan exerts a more pronounced effect on chain configuration after rearrangement.

**FIG. 4.**
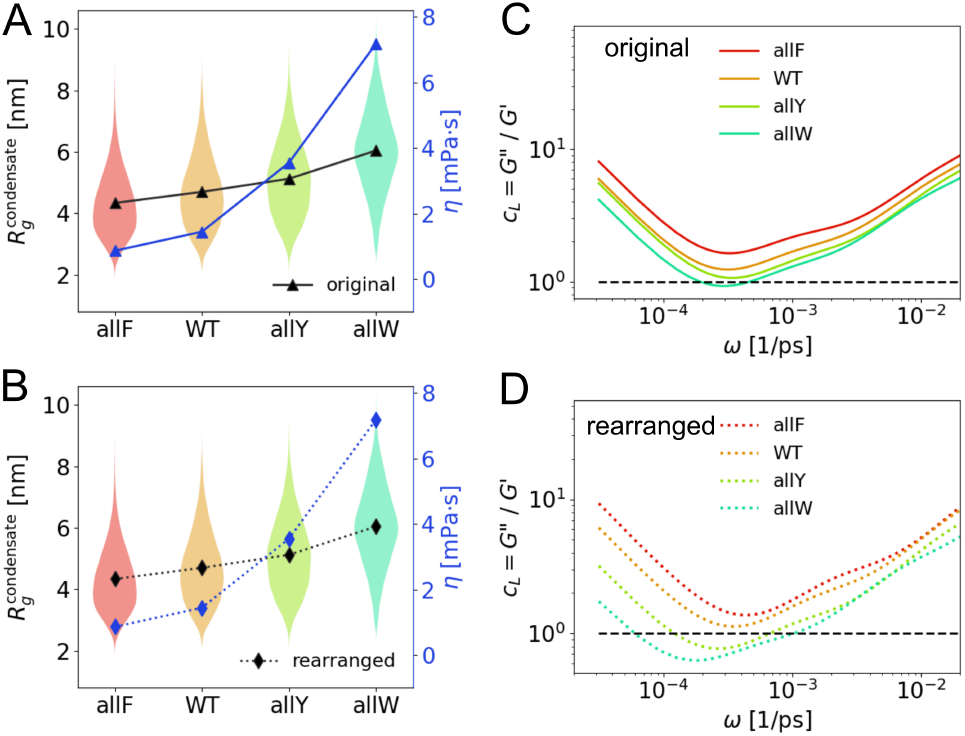
Comparison between original (A and C) and rearranged (B and D) sequences. **(A)** Radius of gyration *R*_*g*_ and viscosity *η* are measured from condensate simulations of four aromatic mutants before rearrangement (triangles and solid lines). **(B)** *R*_*g*_ and *η* are measured from condensate simulations after rearrangement type II (diamonds and dashed lines). **(C-D)** Comparison of *c*_*L*_(*ω*) from condensate simulations between before (solid lines) and after (dashed lines) rearrangement type II. All results come from parameter setting *k* = 16, *θ*_0_ = 140^°^, and residue-dependent angle scheme II.

Next, the allW sequence shows substantial changes in viscoelastic properties upon rearrangement, with viscosity (*η*) increasing by 170% and the minimum loss factor 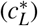 decreasing by 40%. In contrast, the allF sequence shows only modest changes: 14% increase in *η* and 10% decrease in 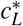 (Fig. 4). These changes from allF to allW in moduli are also consistent with the mutant trends observed earlier (Fig. 3E–F).

Together, these results demonstrate that both the type of aromatic residue (correlating with hydrophobicity and local rigidity) and their spatial concentration enhance the viscous and elastic moduli, promoting a more elastic-dominated regime. Furthermore, the strength and density of aromatic residues act cooperatively to modulate condensate viscoelasticity.

To reveal how aromatic clustering alters condensates’ internal structural and dynamic heterogeneity, we visualized condensates according to the hydrophobicity scale *λ* (Fig. 5A and C). We also computed the self-part of the van Hove correlation function *G*_*s*_(*r, t*). The self-part of the van Hove correlation function, *G*_*s*_(*r, t*) is scaled by 4*πr*^2^, as a function of the radial displacement *r/σ*, where *σ* is a characteristic molecular length scale. The function *G*_*s*_(*r, t*) describes the probability density that a particle has moved a distance *r* after a time interval *t*, and serves as a key indicator of spatial and temporal dynamic heterogeneity in the system. In ideal diffusive systems, *G*_*s*_(*r, t*) is expected to follow a Gaussian distribution, whereas deviations from Gaussianity indicate the presence of dynamic heterogeneity.

**FIG. 5.**
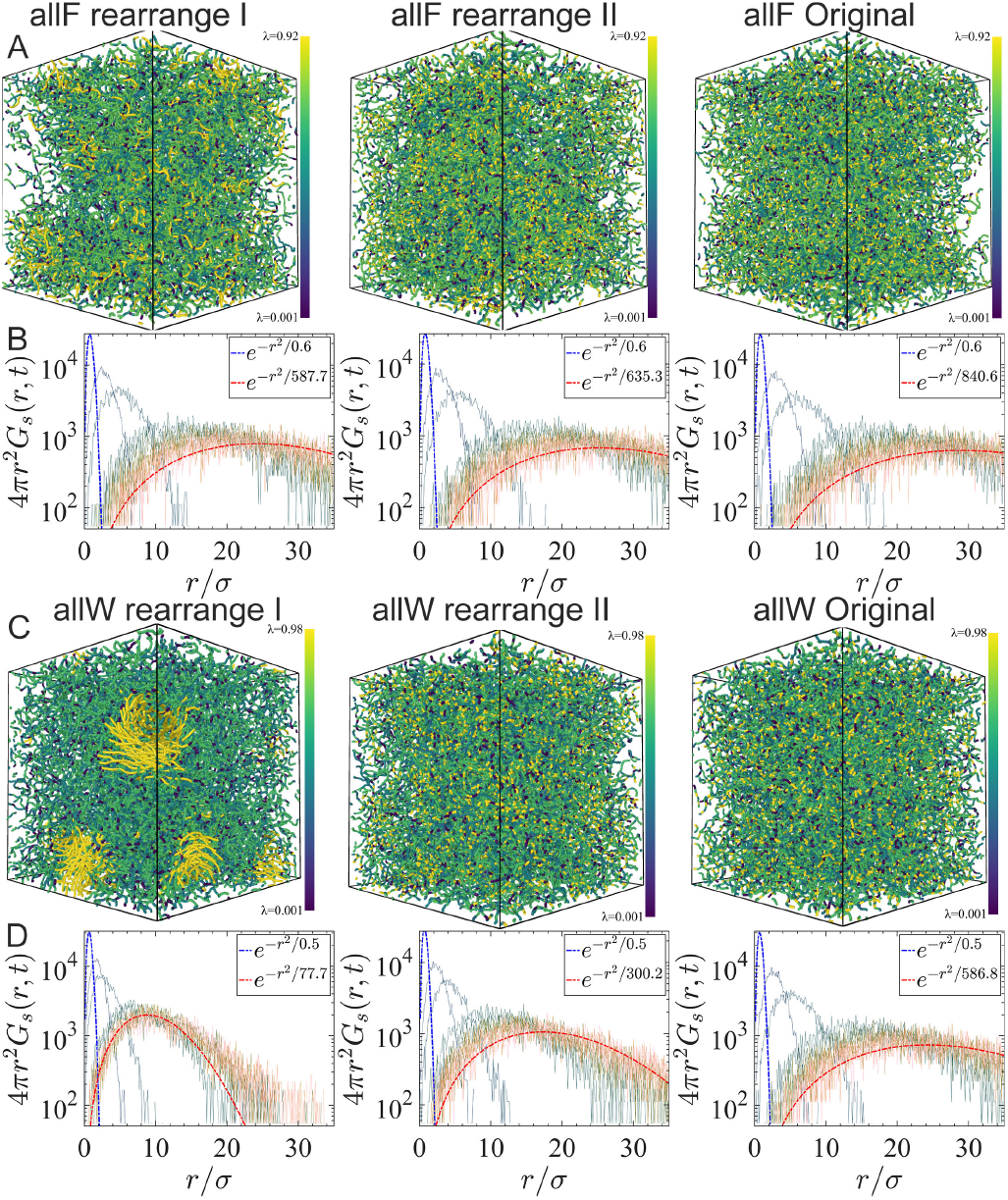
Structural analysis of aromatic mutant systems and their rearranged sequences. **(A and C)** Three-dimensional visualizations of the spatial distribution of molecules colored by the increasing hydrophobicity scale *λ*. Panels show the rearrangement I (left), rearrangement II (center), and original configuration (right), for the allF system (A) and allW system (C). **(B and D)** Corresponding self vanHove functions 4*πr*^2^*G*_*s*_(*r, t*) plotted as a function of distance *r/σ* for each configuration. The Gaussian decay fit parameter, 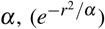 are indicated in each panel. The blue and red fitting curves correspond to *t* = 0.5 ps and *t* = 1000 ps cases.

In each panel (Fig. 5B and D) the numerical data are compared to a Gaussian-like exponential decay of the form *e*^−*r*2*/α*^. The fit parameter *α* characterizes the width of the distribution and is observed to increase with increasing structural randomness. This behavior indicates that molecular rearrangement (from left to right in Fig. 5) leads to more spatially extended displacement distributions and, consequently, to a greater degree of dynamic delocalization. Configurations with larger *α* values correspond to systems with greater spatial disorder and broader dynamic heterogeneity. Comparisons among rearrangement I, II, and the original structures reveal systematic spatial organization and dynamical behavior differences. In the allW rearrangement I system (panel C, left), clusters of high-*λ* residues become more prominent and spatially segregated, with the most attractive residues forming tight clusters. This spatial organization correlates with a sharper deviation from Gaussian behavior and a smaller *α* value in the *G*_*s*_(*r, t*) distribution. In contrast, the original configurations display more homogeneous potential depth distributions and higher *α* values, indicating less spatial heterogeneity and more Gaussian-like dynamics.

### Viscoelastic universality in phase-separated IDP condensates

Beyond uncovering how sequence modulates condensate properties, our findings reveal a broader relationship between chain conformation and viscoelastic behavior invariant to parameterization and sequences. By appropriately rescaling the viscoelastic moduli, we uncover unifying trends that collapse diverse simulation outcomes onto master curves. This indicates that within the coarse-grained energy functions we considered for IDPs, a single dominant relaxation mechanism controls both the elastic and viscous responses across different sequences.

We analyzed data across four categories of parameter settings: (1) variations in the preferred bond angle *θ*_0_, (2) changes in the angular stiffness constant *k*, (3) introduction of residue-dependent angular potentials 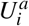, and (4) rearrangements of aromatic sequence patterns (see Table S1, SI-A). Despite the heterogeneity in input parameters and sequences considered, a consistent dynamic response emerges: the loss factor *c*_*L*_(*ω*), which quantifies the ratio of viscous to elastic contributions and exhibits a well-defined minimum at a characteristic frequency *ω*^*\**^. When normalized by this minimum value 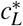 and frequency *ω*^*\**^, all data collapse onto a single curve in the low-frequency regime (Fig. 6A). This collapse reveals that each condensate’s viscoelastic profile can be effectively characterized by a single descriptor, 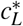, marking the point of maximal elasticity.

**FIG. 6.**
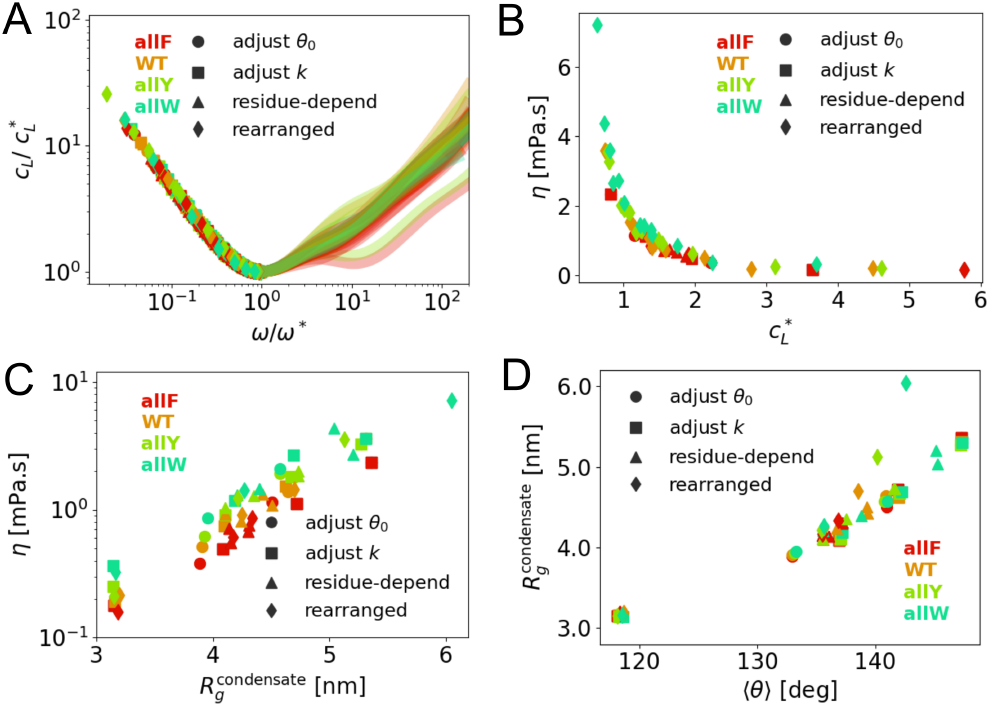
General relations of viscoelasticity and chain configuration. All following panels contain four data sets (see Table S1) and four mutants. The legends “adjust *θ*_0_ (circle)”, “adjust *k* (squares)”, “residue-depend (triangles)”, and “rearranged (diamonds)” represent four categories of data sets in Table S1, respectively. Four mutants of A1-LCD are represented in different colors. **(A)** Normalized loss factor 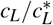 as a function of normalized frequency *ω/ω*^*\**^. The low-*ω* part of all data follows the same curve. **(B)** Viscosity *η* as a function of characteristic loss factor 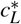.All data collapses into a master curve. **(C)** *η* as a function of *R*_*g*_ in condensates. **(D)** *R*_*g*_ from condensate simulations as a function of average bond angle ⟨*θ*⟩.

Strikingly, we also observe a robust inverse correlation between the shear viscosity *η* and 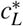 (Fig. 6B): systems with lower 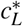 exhibit higher viscosities and stronger elastic dominance. A similar trend is observed in the experiment data (see Fig. S3 of SI-D). This trend reflects a general principle of polymer rheology, underpinned by the Kramers–Kronig relations, which link dissipative and elastic mechanical responses [49].

Viscosity also scales positively with the radius of gyration *R*_*g*_ of chains within condensates (Fig. 6C). Extended conformations enhance inter-chain interactions and entanglements, contributing to higher viscosity. However, aromatic substitutions (e.g., F → Y → W) can increase viscosity without altering *R*_*g*_, highlighting that interaction strength, beyond chain shape, plays a decisive role in viscoelastic control. The relationship between viscosity and interaction energy has been investigated in Fig. S4A of SI-E. Lastly, we show that *R*_*g*_ increases with the average bond angle ⟨*θ*⟩, a proxy for local backbone rigidity (Fig. 6D for 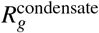 and Fig. S4B for 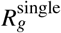). While most data fall along this correlation, rearranged sequences (diamond symbols) deviate, exhibiting greater *R*_*g*_ than expected, indicating that both sequence patterning and rigidity contribute independently to chain expansion and network formation.

The observed collapse of viscosity versus normalized loss factor suggests that a standard underlying timescale governs static and dynamic viscoelastic properties. Indeed, the dynamical behaviour of IDP condensates can be rationalized within sticky Rouse models of associative polymers where transient polymer networks form sticker–sticker interactions and dissociate dynamically [50]. The average lifetime of these reversible crosslinks defines a dominant relaxation timescale *τ*_*R*_, which governs both viscous dissipation and elastic storage. In previous works, we have measured the timescale through the quantification of flow activation energies of condensates in experiments and in simulations [19, 38]. This single timescale behavior can be captured by a single-mode Maxwell model, where the frequency-dependent storage and loss moduli are given by:

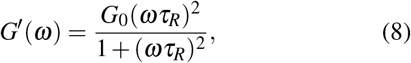

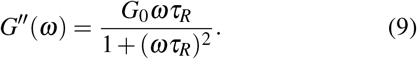

The corresponding loss factor, defined as the ratio of loss to storage modulus, becomes:

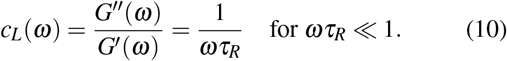

Rescaling *c*_*L*_(*ω*) by its minimum value 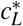 and the corresponding frequency *ω*^*\**^ effectively eliminates differences in *τ*_*R*_ between systems, revealing a shared relaxation mechanism. Variations in chain rigidity or sequence composition alter the value of *τ*_*R*_, but not the functional form of *c*_*L*_(*ω*), leading to the observed collapse onto a universal master curve. Normalizing the loss factor removes sequence-specific timescale differences. It reveals the universal features of the relaxation mechanisms that emerge in IDP condensates within the framework of single-bead, semiflexible coarse-grained models.

## DISCUSSION

Biomolecular condensates exhibit tightly regulated material properties, and their dysregulation is linked to pathological solidification in neurodegenerative diseases [10, 20, 51]. While molecular simulations have advanced our understanding of condensate formation and thermodynamics, standard coarse-grained models that treat IDPs as fully flexible polymers fail to capture their viscoelastic behavior [30, 31]. These models often portray condensates as predominantly viscous, in contrast to experimental microrheology, which reveals a substantial elastic component.

Here, we demonstrate that introducing sequence-dependent backbone rigidity via angular potentials enables coarsegrained simulations to recover key viscoelastic signatures observed in hnRNP A1-LCD condensates [39]. Increasing local stiffness expands the frequency range dominated by elasticity and shifts the viscoelastic crossover, a feature absent in flexible models. Furthermore, guided by the recently proposed energy landscape model of condensate viscoelasticity [31], we showed that rearranging aromatic residues to enhance sticker clustering amplifies these effects, highlighting the role of sequence patterning in regulating condensate material properties.

Beyond reproducing experimental trends, our results reveal a broader organizing principle: a sequence-independent minimum in the loss factor correlating inversely with shear viscosity across diverse parameter settings. This universal behavior suggests that condensate viscoelasticity is governed by a common relaxation mechanism rooted in the transient connectivity of sticker-mediated networks [19, 38]. By normalizing the loss factor and frequency axis, we collapse diverse systems onto a single master curve, exposing a shared viscoelastic response that emerges from the interplay of chain rigidity and interaction dynamics.

Together, these findings establish a general framework linking sequence-encoded chain stiffness to the emergent material properties of protein condensates. Future experiments, such as optical tweezers-based microrheology and multi-scale simulations, could further test and refine this connection, advancing our ability to predict and program condensate mechanics from the sequences and composition of biomolecules.

## ACKNOWLEDGEMENTS

This work was supported by funds from the National Institute of General Medical Sciences, with grant no. R35 GM138243 awarded to DAP.

