## Supporting Information for "Backbone Rigidity Encodes Universal Viscoelastic Signatures in Biomolecular Condensates"

#### A. Detailed sequence and simulation parameter settings

All simulations were performed at $T = 300K$ and the number of A1-LCD chains is 250. We set the equilibrium density of condensates as the same as the condensed phase in phase separation simulations. After calculation, the box lengths of sequence "allF, WT, allY, allW" are $L$= 23.54, 22.75, 22.16, 21.74 nm, respectively. We gradually shrinking the box dimensions from 50 nm to the target size, increasing the density of our system until it reaches the condensate density in previous phase separation simulations

The detailed amino acid sequence of four aromatic mutants of A1-LCD and their rearrangements are listed in Table S1. The aromatic residues in WT has 12 F and 8 Y residues. They are replaced to "F, Y and W" in "allF, allY and allW" sequences, respectively.

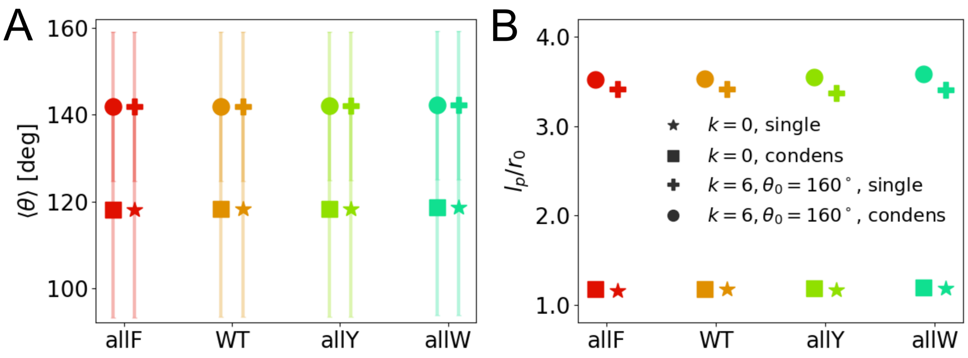


Figure S1: A. Average bond angle $\langle\theta\rangle$ for four mutants of A1-LCD (allF, WT, allY, allW) under two different $U^a$ parameter sets. Squares and circles indicate parameters $k = 0$ and $k = 6\ k_BT/\mathrm{rad}^2$, $\theta_0 = 160°$ in condensate simulations. Stars and crosses indicate them in single-chain simulations. Errorbars show standard deviation of $\theta$ distribution. B. Persistence length $l_p$ for four mutants. The simulation conditions are the same as panel A.

### B. Quantification of local stiffness and persistence length

We quantify the local stiffness with distribution of bond angle $\theta$. Taking all residues in the system (single-chain or condensates) as samples, we calculated average angle $\langle\theta\rangle$ and standard deviation (errorbar) as shown in Fig. S1.

After the angular potential ($k = 6$, $\theta_0 = 160°$) is applied, $\langle\theta\rangle$ increases from 118° to 142°. We also observed that ($\langle\theta\rangle = 142°$) is slightly lower than the natural angle setting ($\theta_0 = 160°$) in angular potential, due to other non-bonded interactions, e.g. steric interaction, among residues. Notably, $\langle\theta\rangle$ remains consistent between single-chain and condensate simulations across different A1-LCD mutants (sequences allF, WT, allY and allW have increasing hydrophobicity).

#### Persistence length

Since bond length remains constant ($r_0 = 0.38$ nm) for all residues, we calculated the persistence length as $l_p/r_0 = -1/\ln\langle\cos(\pi - \theta)\rangle$, which serves as an alternative measure of local stiffness. In all our simulations, $l_p$ is less than 50% of gyration radius $R_g$, indicating that the peptide remains flexible even under the applied angular potential.

### C. Modulation of viscoelastic moduli

As local stiffness and hydrophobic interactions increases, viscosity $\eta$ also rises due to stronger inter-chain interactions and potential entanglement. The corresponding viscosity values of simulations in Fig. 2 are shown in Fig. S2A.

Viscoelastic moduli respond to variations in torsional stiffness $k$ and the angular potential's natural angle $\theta_0$. When $k$ increases from 0 to 20 ($\theta_0 = 160°$), local stiffness strengthens and the chain straightens. Consequently, $G'$ grows more rapidly than $G''$, leading to the emergence of a dominantly elastic regime (Fig. S2B). Similarly, increasing $\theta_0$ from 140° to 160° at $k = 10$ induces a comparable transition between viscous and elastic regimes (Fig. S2C).

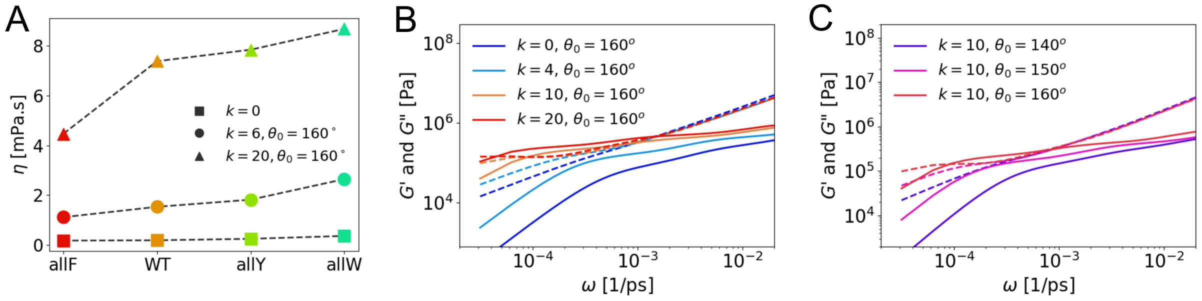


Figure S2: A. Viscosity $\eta$ measured from four mutants simulations at three parameters $k = 0$; $k = 6$, $\theta_0 = 160°$; and $k = 20$, $\theta_0 = 160°$. B. $G'(\omega)$ (solid lines), $G''(\omega)$ (dashed lines) for WT mutant at varying $k$ parameters. C. $G'(\omega)$, $G''(\omega)$ for WT mutant at varying $\theta_0$ parameters.

### D. Robust viscoelastic correlation in experiment

In Fig. 1g of,[1] the loss factor $c_L(\omega)$ consistently exhibits a characteristic dynamic behavior: a turning point $c_L^*$, marking a transition from a plateau to a decay at frequency $\omega^*$. After normalization data across different mutants, we observed a similar collapse, as shown in Fig. S3A. An inverse correlation between the viscosity $\eta$ and $c_L^*$ was also observed in Fig. S3B.

Although various factors in the experimental system—such as hydrodynamic effects—may introduce fluctuations in this relationship, the persistence of this correlation suggests that

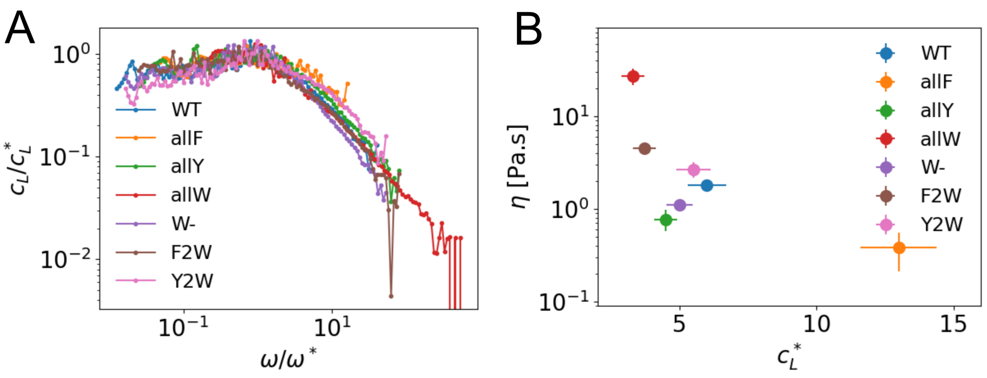


Figure S3: A. Normalized loss factor $c_L/c_L^*$ as a function of normalized frequency $\omega/\omega^*$. B. Viscosity $\eta$ as a function of characteristic loss factor $c_L^*$. All data comes from the experiment[1] at temperature 26°C.

universal features of the underlying relaxation mechanisms remain detectable in realistic systems.

### E. Other general relationships

#### Characteristic coarse-grained energy

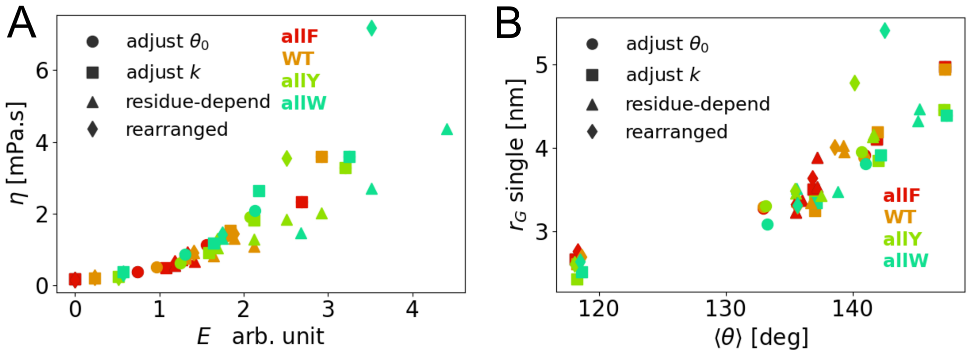


Figure S4: A. Viscosity $\eta$ as a function of energy $E = E^a + 100E^H$. The lowest energy point has been set to 0 for convenience. B. $R_g$ from single-chain simulations as a function of average bond angle $\langle\theta\rangle$.

In previous study, the attractive energy strength is found to be related to the viscoelasticity. From our observation, we consider to add the angular energy strength which relates to the chain extension. Here, we observe the relationship between viscosity $\eta$ and coarse-grained energy $E$, which is a linear superposition of angular potential part ($E^a$) and hydrophobic interaction part ($E^H$):

$$E = E^a + c_E E^H. \tag{S1}$$

The electrostatic interaction part is not included because the charged residues keep the same in our simulations. The angular potential part $E^a = \frac{k}{2}\langle(\theta_{0,i} - \theta^*)^2\rangle_i$, where $i$ is the residue index and $\theta^* = 118°$ is the equilibrium bond angle without any angular potential, represents the characteristic angular potential of a sequence. The hydrophobic interaction part $E^H = \epsilon\langle\lambda_i\rangle_i$, represents the attractive depth of residue hydrophobic potential. We estimate the coefficient $c_E$ by comparing the viscoelasticity between different angular potential settings and mutants. As an example, we take several data points at $\eta \approx 1.8$ mPa·s on Fig. 5B, and calculate their Characteristic energies $E^a$ and $E^H$. The similar viscosity values should be related to similar energies. From this comparison, we get $c_E \approx 100$ and generalize this coefficient for our whole data settings, as shown in Fig. S4A.

#### Single-chain $R_g$

Similar to condensed simulations, radius of gyration $R_g$ in single-chain simulation is also positively correlated with local stiffness represented by $\langle\theta\rangle$ (Fig. S4B). Because the intra-chain interactions from varying hydrophobicity of different sequences would modify $R_g$ more than the condensed cases, more fluctuations appear in this general relationship.

Table S1: The detailed amino acid sequence of four aromatic mutants of A1-LCD and their rearrangements

| Name | Amino acid sequence |
|---|---|
| allF | GSMASASSSQRGRSGSGNFGGGRGGGFGGNDNFGRGGNFSGRGGFG<br>GSRGGGGFGGSGDGFNGFGNDGSNFGGGGSFNDFGNFNNQSSNFGP<br>MKGGNFGGRSSGGSGGGGQFFAKPRNQGGFGGSSSSSSFGSGRRF |
| WT | GSMASASSSQRGRSGSGNFGGGRGGGFGGNDNFGRGGNFSGRGGFG<br>GSRGGGGYGGSGDGYNGFGNDGSNFGGGGSYNDFGNYNNQSSNFGP<br>MKGGNFGGRSSGPYGGGGQYFAKPRNQGGYGGSSSSSSYGSGRRF |
| allY | GSMASASSSQRGRSGSGNYGGGRGGGYGGNDNYGRGGNYSGRGGYG<br>GSRGGGGYGGSGDGYNGYGNDGSNYGGGGSYNDYGNYNNQSSNYGP<br>MKGGNYGGRSSGGSGGGGQYYAKPRNQGGYGGSSSSSSYGSGRRY |
| allW | GSMASASSSQRGRSGSGNWGGGRGGGWGGNDNWGRGGNWSGRGG<br>WGGSRGGGGWGGSGDGWNGWGNDGSNWGGGGSWNDWGNWNN<br>QSSNWGPMKGGNWGGRSSGSGGGGGQWWAKPRNQGGWGGSSSS<br>SSWGSGRRW |
| Rearranged (II), allF | GSMASASSSQRGRSGSGNGGGRGGGGGNDNGRGGNSGRGGGGSRGG<br>GGGFGFSFGFDFGFNFGFGFNFDFGFSFNFGFGFGFGFSFNDGNNN<br>QSSNGPMKGGNGGRSSGGSGGGGQAKPRNQGGGGSSSSSSGSGRR |
| Rearranged (II), WT | GSMASASSSQRGRSGSGNGGGRGGGGGNDNGRGGNSGRGGGGSRGG<br>GGGFGFSFGFDFGYNYGFGFNYDFGYSFNFGYGYGFGYSYNFDGNNN<br>QSSNGPMKGGNGGRSSGPGGGGQAKPRNQGGGGSSSSSSGSGRR |
| Rearranged (II), allY | GSMASASSSQRGRSGSGNGGGRGGGGGNDNGRGGNSGRGGGGSRGG<br>GGGYGYSYGYDYGYNYGYGYNYDYGYSYNYGYGYGYGYSYNDGNN<br>NQSSNGPMKGGNGGRSSGGSGGGGQAKPRNQGGGGSSSSSSGSGRR |
| Rearranged (II), allW | GSMASASSSQRGRSGSGNGGGRGGGGGNDNGRGGNSGRGGGGSRGG<br>GGGWGWSWGWDWGWNWGWGWNWDWGWSWNWGWGWGWGW<br>SWNDGNNNQSSNGPMKGGNGGRSSGSGGGGGQAKPRNQGGGGSS<br>SSSSGSGRR |

Table S2: The parameter lists of various torsional angular potential $U^a$: stiffness $k$ and natural angle $\theta_{0,i}$. The data sets $U^a$-$\theta_0$ and $U^a$-$k$ have constant natural angle $\theta_0$. The other two data sets $U^a$-$\theta_{\mathrm{Ar}}$ and Re have residue-dependent $\theta_{0,i}$, which is represented by $\theta_{\mathrm{Ar}}$ and $\theta_{\mathrm{Ar}\pm}$ ("Ar" represents aromatic amino acid F,Y or W).

| Data set | $k$ $[k_BT/\mathrm{rad}^2]$ | $\theta_0$ [deg] | $\theta_{\mathrm{Ar}}$ [deg] | $\theta_{\mathrm{Ar}\pm}$ [deg] |
|---|---|---|---|---|
| adjust $\theta_0$ | 10 | 140 | $\theta_{\mathrm{Ar}} = \theta_0$ | $\theta_{\mathrm{Ar}\pm} = \theta_0$ |
| adjust $\theta_0$ | 10 | 150 | $\theta_{\mathrm{Ar}} = \theta_0$ | $\theta_{\mathrm{Ar}\pm} = \theta_0$ |
| adjust $\theta_0$ | 10 | 160 | $\theta_{\mathrm{Ar}} = \theta_0$ | $\theta_{\mathrm{Ar}\pm} = \theta_0$ |
| adjust $k$ | 0 | / | / | / |
| adjust $k$ | 4 | 160 | $\theta_{\mathrm{Ar}} = \theta_0$ | $\theta_{\mathrm{Ar}\pm} = \theta_0$ |
| adjust $k$ | 6 | 160 | $\theta_{\mathrm{Ar}} = \theta_0$ | $\theta_{\mathrm{Ar}\pm} = \theta_0$ |
| adjust $k$ | 10 | 160 | $\theta_{\mathrm{Ar}} = \theta_0$ | $\theta_{\mathrm{Ar}\pm} = \theta_0$ |
| adjust $k$ | 20 | 160 | $\theta_{\mathrm{Ar}} = \theta_0$ | $\theta_{\mathrm{Ar}\pm} = \theta_0$ |
| residue-depend (0) | 16 | 140 | $\theta_{\mathrm{Ar}} = \theta_0$ | $\theta_{\mathrm{Ar}\pm} = \theta_0$ |
| residue-depend (I) | 16 | 140 | $\theta_{\mathrm{F}} = 145$, $\theta_{\mathrm{Y}} = 160$, $\theta_{\mathrm{W}} = 175$ | $\theta_{\mathrm{Ar}\pm} = \theta_0$ |
| residue-depend (II) | 16 | 140 | $\theta_{\mathrm{F}} = 145$, $\theta_{\mathrm{Y}} = 160$, $\theta_{\mathrm{W}} = 175$ | $\theta_{\mathrm{Ar}+} = \theta_{\mathrm{Ar}}$, $\theta_{\mathrm{Ar}-} = \theta_0$ |
| residue-depend (III) | 16 | 140 | $\theta_{\mathrm{F}} = 145$, $\theta_{\mathrm{Y}} = 160$, $\theta_{\mathrm{W}} = 175$ | $\theta_{\mathrm{Ar}\pm} = \theta_{\mathrm{Ar}}$ |
| Rearranged (0) | 0 | / | / | / |
| Rearranged (I) | 16 | 140 | $\theta_{\mathrm{Ar}} = \theta_0$ | $\theta_{\mathrm{Ar}\pm} = \theta_0$ |
| Rearranged (II) | 16 | 140 | $\theta_{\mathrm{F}} = 145$, $\theta_{\mathrm{Y}} = 160$, $\theta_{\mathrm{W}} = 175$ | $\theta_{\mathrm{Ar}+} = \theta_{\mathrm{Ar}}$, $\theta_{\mathrm{Ar}-} = \theta_0$ |
